# KASSPer: Kinase Active Site Structure Prediction using Protein and Ligand Language Models and Its Application to Virtual Screening

**DOI:** 10.64898/2026.01.07.698135

**Authors:** Wonkyeong Jang, Woong-Hee Shin

**Affiliations:** Department of Biomedical Informatics, Korea University College of Medicine, Seoul 02708, Republic of Korea; Arontier. Co., Seoul 06735, Republic of Korea

## Abstract

**Motivation:** Structure-based virtual screening (SBVS) is limited by the rigid-receptor assumption, which is particularly problematic for kinases that adopt multiple active-site conformations but are experimentally biased toward a single state. Although ensemble screening can address this limitation, it remains computationally expensive.

**Results:** We introduce KASSPer (Kinase Active Site Structure Predictor), a framework that predicts kinase active-site conformational states using protein and compound language models. Given a kinase amino acid sequence and a ligand SMILES string, KASSPer enables ligand-specific conformer selection prior to SBVS, substantially reducing the computational cost associated with ensemble screening. Benchmarking on the DUD-E kinase subset demonstrates that KASSPer-guided screening consistently outperforms ensemble-based approaches across all evaluation metrics.

**Availability and Implementation:** The implementation for model loading and inference is available at the GitHub repository https://github.com/kucm-lsbi/KASSPer

## 1. Introduction

Structure-based virtual screening (SBVS) has become an indispensable part of drug discovery by utilizing the three-dimensional structural information of target proteins to predict protein–ligand binding poses and corresponding affinities to find promising hit candidates from a chemical library. Despite its proven utility, SBVS often struggles to account for receptor conformational flexibility. Structural states and binding-site geometries can vary substantially during ligand binding (Bordogna *et al*. 2011, Fan and Irwin 2009). This limitation can lead to false positives when screening against a single static structure, thereby reducing hit rates and downstream success in hit identification.

To address this challenge, ensemble screening employs multiple receptor conformations derived from X-ray crystallography, NMR ensembles, or molecular dynamics simulations to capture the conformational space of the target diversely (Modi and Dunbrack 2019). In this approach, a given ligand is docked to multiple receptor structures. A representative score is then selected for compound ranking. While ensemble screening can improve the finding of hit molecules from the library, it comes at the expense of substantial computational cost, as each ligand must be docked across an entire set of receptor structures. Furthermore, selecting representative conformations from a large pool remains nontrivial and may introduce bias if the structural ensemble is not sufficiently diverse or balanced.

Protein kinases represent one of the most extensively studied and therapeutically relevant target families, with over 500 members regulating key cellular processes. Kinase active sites are characterized by the conformations of the DFG motif and the αC-helix. These structural features give rise to well-defined binding modes, including DFG-in, DFG-out, and intermediate states (Berman *et al*. 2000). Tools like KinCoRe classify kinase conformations by analyzing activation-loop positions, αC-helix orientation, and DFG dihedral angles (Modi and Dunbrack 2022). However, the Protein Data Bank (PDB) remains heavily biased toward the DFG-in BLAminus state, limiting the discovery of inhibitors that preferentially bind less-populated or transient kinase conformations.

Recent advances in the protein structure prediction field, such as AlphaFold (Abramson *et al*. 2024, Krishna *et al*. 2024, Passaro *et al*. 2025) and its derivatives, have demonstrated remarkable improvements in accuracy and have been applied to generate diverse conformations for ensemble screening workflows (Song *et al*. 2024). Despite these advancements, docking molecule library against large numbers of predicted models requires high computational demand and may yield redundant or non-physiological conformations.

In this study, we introduce KASSPer (<u>K</u>inase <u>A</u>ctive <u>S</u>ite <u>S</u>tructure <u>P</u>r<u>e</u>dictor). This novel algorithm predicts the most probable kinase conformation adopted upon binding of a given ligand. The program utilizes the primary sequence of the kinase and ligand descriptors as inputs. To develop a predictive framework for kinase–ligand compatibility, a range of traditional machine-learning algorithms and deep learning architectures were evaluated to ensure robust performance across various modeling approaches. In accordance with the benchmark study, a stacking ensemble combining LightGBM and Random Forest (RF) was selected as the predictor. By predicting a receptor conformation tailored to each query compound, KASSPer restructures the virtual screening pipeline. Instead of exhaustive ensemble screening, KASSPer adopts a targeted strategy in which each compound is docked to its predicted structure. We benchmarked this approach using the DUD-E (Mysinger *et al*. 2012) kinase subset. The study revealed substantial enhancements in enrichment factors at early retrieval rates, thereby underscoring its capacity to expedite the identification of conformation-selective kinase inhibitors. This strategy has the potential to enhance the efficacy of SBVS, facilitating the identification of hit compounds for various targets with structural diversity.

## 2. Materials and Methods

### 2.1 Benchmark Dataset for Kinase Structure Prediction and Annotation

We extracted human kinase crystal structures from KLIFS database (Kanev *et al*. 2020) (accessed May 2025). The database contains catalytic domain structures of kinases, extracted from the Protein Data Bank (PDB), as well as their respective inhibitors. In addition, it provides interaction information between the protein and the compound. Out of 6336 human crystal structures with bound ligand in KLIFS, 50 PDBs could not be accessed, resulting in 6286 structures. After splitting PDB structures into chains and removing redundant protein chain-ligand, 7673 pairs remained.

The conformational states of the benchmark set were annotated by Kinase Conformation Resource (KinCoRe) (Modi and Dunbrack 2022) standalone version. The binding site of kinases contains an activation loop that features the Asp-Phe-Gly (DFG) motif in the N-terminal region. This motif plays a crucial role in anchoring ATP to the active site. KinCoRe initially classifies kinase conformations based on the position of the activation loop. The three primary states are DFGin, DFGinter, and DFGout. In the DFGin state, Phe of the motif is located within the ATP-binding pocket, enabling the kinase to bind ATP. This configuration is designated as the active state. Conversely, DFGout conformation directs the Phe out of the ATP binding pocket, thus classifying it as an inactive state. The three primary classes are subdivided according to the backbone dihedral angles of X-DF and the χ1 angle of Phe, where X denotes a residue preceding the motif. In the system, the letters A, B, and L are assigned to alpha, beta, and left-handed, respectively. The Phe χ1 angle is designated as plus, minus, and trans for π/3, -π/3, and π, respectively. Consequently, the predominant class, designated as DFGin, comprises seven distinct subclasses: BLAminus, BLAplus, ABAminus, BLBminus, BLBplus, BLBtrans, and Unassigned. In contrast, DFGinter (comprising BABtrans and Unassigned) and DFGout (consisting of BBAminus and Unassigned) are characterized by a mere two subclasses. In the present study, since the sample size for subclasses is inadequate, the two subclasses of DFGout and DFGinter with Unassigned_Unassigned were aggregated and denoted as the ’DFGout’ and ‘DFGothers’ classes, respectively. Ultimately, nine structural states—ABAminus, BLAminus, BLAplus, BLBminus, BLBplus, BLBtrans, Unassigned, DFGothers, and DFGout—were designated as the prediction targets of the model.

The human kinase benchmark dataset was divided into three sets: a training set comprising 70% of the total data, a validation set (15%), and a test set (15%) using a stratified split. Specifically, we utilized the stratify option of StratifiedShuffleSplit and train_test_split from scikit-learn (Pedregosa *et al*. 2011) to divide the dataset, under the assumption that the distribution of classes in each set was equivalent to the overall dataset.

To validate the cross-species generalization, 396 mouse kinase structures were also extracted from the KLIFS database. After removing duplicated ligands, 303 unique protein-ligand pairs were obtained and constructed into the mouse benchmark set. The dataset was constructed using the same protocol as the human kinase. In contrast to human kinases, none of the structures were annotated as BLBtrans, resulting in eight states. The distribution of crystal structures for each class is shown in Supplementary data Figure S1.

The sequence similarities between the human kinase test set and the mouse set were also analyzed using MMSeqs2 (Version 17.b804f) to measure how similar they are. Pairwise sequence comparisons were conducted using the search module, with a sensitivity of 7.5, an E-value cutoff of 1×10LL, and coverage mode 2, with a minimum coverage threshold of 0.5. Also, the compound similarities were measured using ECFP4-based Tanimoto similarity. The distribution of pairwise sequence and compound similarities is illustrated in Supplementary data Figure S2.

### 2.2 Pretrained Language Models for Kinases and Compounds

To predict the conformational state of a kinase for a given compound, we utilized language models to embed molecules. To extract protein sequence embeddings, we used the ESM2 (Lin *et al*. 2023) protein language model. For each protein sequence, we utilized the output of the last hidden state of the ESM2 model. Only the hidden state values of the valid tokens corresponding to the actual protein sequence, excluding padding tokens, were converted to a fixed-size vector representation by mean pooling. The resulting embeddings were used for subsequent model training.

The SMILES sequence embeddings of the ligands were generated using the ChemBERTa (Chithranada *et al*. 2020) model. ChemBERTa is based on the RoBERTa architecture and is a pre-trained model with a masked language modeling approach using the ZINC dataset. For each ligand’s SMILES sequence, the model’s last hidden state was extracted and converted into a fixed-size vector by performing mean pooling of the sequence tokens. This vector representation was then utilized as input data for model training.

### 2.3 Machine Learning and Deep Learning Models and Hyperparameter Optimization

#### 2.3.1 Machine Learning Models

In this study, we employed a variety of machine learning methods, including tree-based ensemble models, linear models, and support vector machines (SVMs) (Cortes 1995). For tree-based methods, XGBoost (Chen and Guestrin 2016), LightGBM (Ke *et al*. 2017), CatBoost (Prokorenkova *et al*. 2018), and RF (Breiman 2001) were employed. For linear models, the implementation of logistic regression (Cox 1958) and SVM was undertaken. To optimize the hyperparameters of each model, the grid search (Pedregosa *et al*. 2011) was applied to identify the optimal combination. The hyperparameters are listed in the Supplementary data Table S1. All the machine learning models were utilized from scikit-learn.

#### 2.3.2 Deep Learning Models

Two architectures for utilizing deep learning were designed. The first model is concatenation. The embeddings of the protein and ligand were concatenated, and a multi-layer perceptron (MLP) (Rumelhart *et al*. 1986) was constructed to create an input vector. Subsequently, the vector traversed successive layers of hidden nodes, with a depth of 1024, 512, and 256 nodes, respectively. Each hidden layer implemented the rectified linear unit activation function (Nair and Hinton 2010), with a dropout rate of 0.3 applied to mitigate the risk of overfitting.

The second is a cross-attention (Vaswani *et al*. 2017) model that integrates protein and ligand embeddings. Each input embedding was projected to the same dimension through a linear transformation, and then information was exchanged with each other through a bidirectional cross-attention mechanism. The two vectors obtained through the attention mechanism were combined and fed into an MLP for subsequent classification.

Both models were trained following the common protocol. Models were trained for 50 epochs with an initial learning rate of 1e-3. The learning rate was gradually decreased using a cosine annealing scheduler (Loshchilov and Hutter 2016). Adam (Kingma and Ba 2014) and AdamW (Loshchilov and Hutter 2017) were utilized as optimizers, with weight decay exploring values of 1e-6, 1e-5, and 1e-4, and the optimal model was identified based on the Matthews correlation coefficient (MCC) on the validation set.

#### 2.3.3 Stacking Ensemble

To enhance the model’s performance and broaden its generalizability, we employed the stacking ensemble technique (Wolpert 2017), which integrates two models from the top three machine learning models: XGBoost, LightGBM, and RF. From each pre-trained base model, the predicted probabilities were organized into new features and provided as input to a meta-model using logistic regression. The meta-model was trained with the following parameters: multi_class=’multinomial,’ solver=’lbfgs,’ and max_iter=200. Following the training process, the model was utilized to make final classification predictions.

### 2.4 Classification Model Performance Evaluation Metrics

To comprehensively evaluate the performance of the classification model, the MCC was used as the primary metric, and precision, recall, and F1-score served as secondary metrics. All metrics are calculated based on the values of true positive (TP), false positive (FP), true negative (TN), and false negative (FN) in the confusion matrix. Precision is the ratio of TPs to all predicted positives (TP + FP), while recall is the percentage of TPs to all actual positives (TP + FN). F1-score is a harmonized mean of precision and recall that evaluates the balance between the two metrics. The MCC, a class imbalance robust metric, includes all elements of the confusion matrix, calculated as follows:

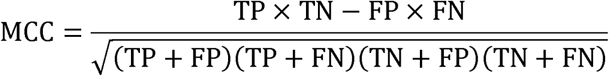

The value of MCC can range from +1 (perfect prediction) to -1 (perfect mismatch).

### 2.5 Applying KASSPer to SBVS and Comparing with Ensemble Screening

#### 2.5.1 Benchmark Dataset

To ascertain the validity of the KASSPer prediction to drug screening, the DUD-E kinase subset, which consists of 26 kinases, was employed. Each kinase possesses a representative structure for SBVS, active compounds that were validated as binders to the target protein, and decoy compounds that do not bind to the receptor but are chemically similar to the actives. Among them, SRC kinases were excluded from the evaluation since the reference crystal structure provided was not of human origin.

With the kinase subset, two benchmarks were performed: docking and screening. For docking, the co-crystallized compound was re-docked to the receptor in order to evaluate the using KASSPer-predicted structures is helpful to reproduce the binding pose. For screening, the active and decoy molecules provided in DUD-E set were docked to the KASSPer-predicted conformation and compared to the ensemble screening provided by Song et al. (Song *et al*. 2024)

#### 2.5.2 KASSPer and Ensemble Screening Protocol

To get the diverse structure of kinases, the structures using AF2 with a multi-state modeling (MSM) protocol (Song *et al*. 2024) were employed. The MSM protocol searches the top five highest sequence-similar templates to the query protein that have the structural state of interest. Instead of generating multiple sequence alignments, MSM starts AF2 from the template structure with an alignment of the sequence pairs to ensure the predicted model has the conformation. After predicting five models per template, KinCoRe is employed to filter out predicted models with the undesired conformation. Among the remained models, the structure with the highest pLDDT is selected for screening.

For the modeled structures, the docking was performed using AutoDock-GPU (Santos-Martins *et al*. 2021). After aligning the model to the crystal structure, a docking box was defined as a cubic box with a side length of 22.5 Å. The center of the box was set to the geometrical center of co-crystallized ligand. AutoDock-GPU was executed with default parameters except for nrun = 50, the number of generated poses. For KASSPer screening, the kinase conformation was predicted for every single compound in the DUD-E set. Subsequently, the molecule was docked to the predicted conformation. In the event that KASSPer predicted a conformation as DFGout, docking was performed for DFGout_BBAminus and DFGout_Unassigned conformations. The result with the lower score was then used for screening. Similarly, in instances where the prediction was designated as DFGothers, the lowest docking score derived from the DFGinter_BABtrans, DFGinter_Unassigned, and Unassigned_Unassigned categories was employed for the screening process. For ensemble screening, the representative pose and score of a compound were selected based on the lowest AutoDock energy across all MSM models.

#### 2.5.3 Evaluation Metrics

The docking benchmark was evaluated by two parameters: the root-mean-square-distance (RMSD) of predicted binding poses to the crystal structure and the docking success, which is defined as RMSD < 2 Å. Spyrmsd (Meli and Biggin 2020), which considers molecular symmetry to calculate RMSD, was utilized to evaluate the metric.

For the VS benchmark, active and decoy molecules were ranked according to the docking score. The area under the curve (AUC) and enrichment factors (EFs) were calculated to evaluate the screening performance. AUC is a metric that quantifies the discriminatory capabilities of a model in distinguishing between active and inactive compounds. AUC values approaching one are indicative of superior performance, and 0.5 corresponds to random retrieval.

EF is an indicator of the ability of an active compound to enrich within a certain percentage of the top docking scores compared to random selection. The enrichment factor at the top x% (EF_x%_) is calculated as follows:

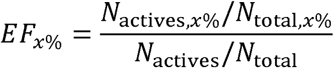

where N_actives,x%_ and N_actives_ are the number of active compounds and in the top x% and the library, respectively. Similarly, N_total,x%_ and N_total_ are the number of active compounds within the top x% and the entire dataset, respectively. In this study, the EF values at the top 1%, 5%, and 10% were calculated.

## 3. Result and Discussion

### 3.1 Performance of KASSPer for Predicting Conformational State

The performance of the prediction models was validated on the human test set (1151 structures) and mouse set (303 structures) to measure cross-species generalization. The human benchmark set was used as the primary evaluation, while the mouse set was employed to provide insight into generalization performance. The top three MCC models on both sets are presented in Table 1. A more detailed result can be found in the Supplementary data Table S2.

**Table 1.** MCC values of the top three single models and stacking ensembles.

| Test Set | Single Model |  |  | Stacking Ensemble |  |  |
| --- | --- | --- | --- | --- | --- | --- |
|  | XGBoost | LightGBM | RF | XGBoost +<br>LightGBM | LightGBM<br>+ RF | XGBoost +<br>RF |
| Human | 0.796 | 0.819 | 0.798 | 0.811 | 0.818 | 0.806 |
| Mouse | 0.602 | 0.554 | 0.580 | 0.563 | 0.558 | 0.586 |

In the human test set, LightGBM attained the highest MCC of 0.819, followed by RF (0.798) and XGBoost (0.796). In contrast, XGBoost demonstrated the highest performance with an MCC of 0.602, followed by RF (0.580) and LightGBM (0.554), on the mouse set. To observe whether stacking ensembles could improve performance and robustness, the top three models were combined. Among the three combinations, the ensemble of LightGBMLandLRF showed the highest MCC values on human data with an MCC of 0.818. The XGBoostLwithLRF ensemble demonstrated the most effective generalization on the mouse data with an MCC of 0.586. All machine learning models exhibited reduced performance on the mouse dataset, with MCC values decreasing by approximately 0.2. This reduction likely reflects fundamental differences between the human and mouse benchmark sets. As illustrated in Supplementary data Figures S1 and S2, the mouse kinases have different conformational distributions and low sequence similarity. The majority of the sequence pairs (71.62%) demonstrate an identity percentage of approximately 20-30%. It is also notable that 92.53% of compound pairs demonstrate a Tanimoto coefficient ranging from zero to 0.2. The low inter-species similarity hampers effective feature learning, yielding lower performance for models trained on mouse kinases compared with those trained on human kinases.

To evaluate the suitability of our model for drug binding conformation prediction, we conducted a comparative analysis focusing on its performance on the human test set. LightGBM and the stacking model of LightGBM and RF, which achieved the highest performance among individual models, were compared. While the stacking model resulted in a marginal decrease in MCC on the human test set (0.001), it achieved a slight improvement on the mouse test set (0.004). Furthermore, the stacking model exhibited a comparable performance to the single LightGBM model in terms of recall, macro precision, and the average macro F1-score.

A more detailed comparison of label-wise prediction performance, as illustrated in Figure 1 and Supplementary data Table S3, revealed that the stacking model led to improved classification for BLAminus, BLBplus, and Unassigned in the mouse dataset. However, a slight performance degradation was observed for DFGout. On the human test set, the performance of the stacking model was slightly reduced due to lower performance for BLBminus, BLBplus, and BLBtrans. In contrast, improved predictions for ABAminus, BLAplus, and Unassigned partially offset these declines. In the case of BLBminus, performance gains observed in the mouse set were accompanied by reduced accuracy in the human test set. This may be due to inherent inter-species variability, suggesting that class-specific patterns learned from one species may not be directly transferable to another. The stacking model combining LightGBM and RF was selected as the final model for KASSPer.

**Figure 1.**
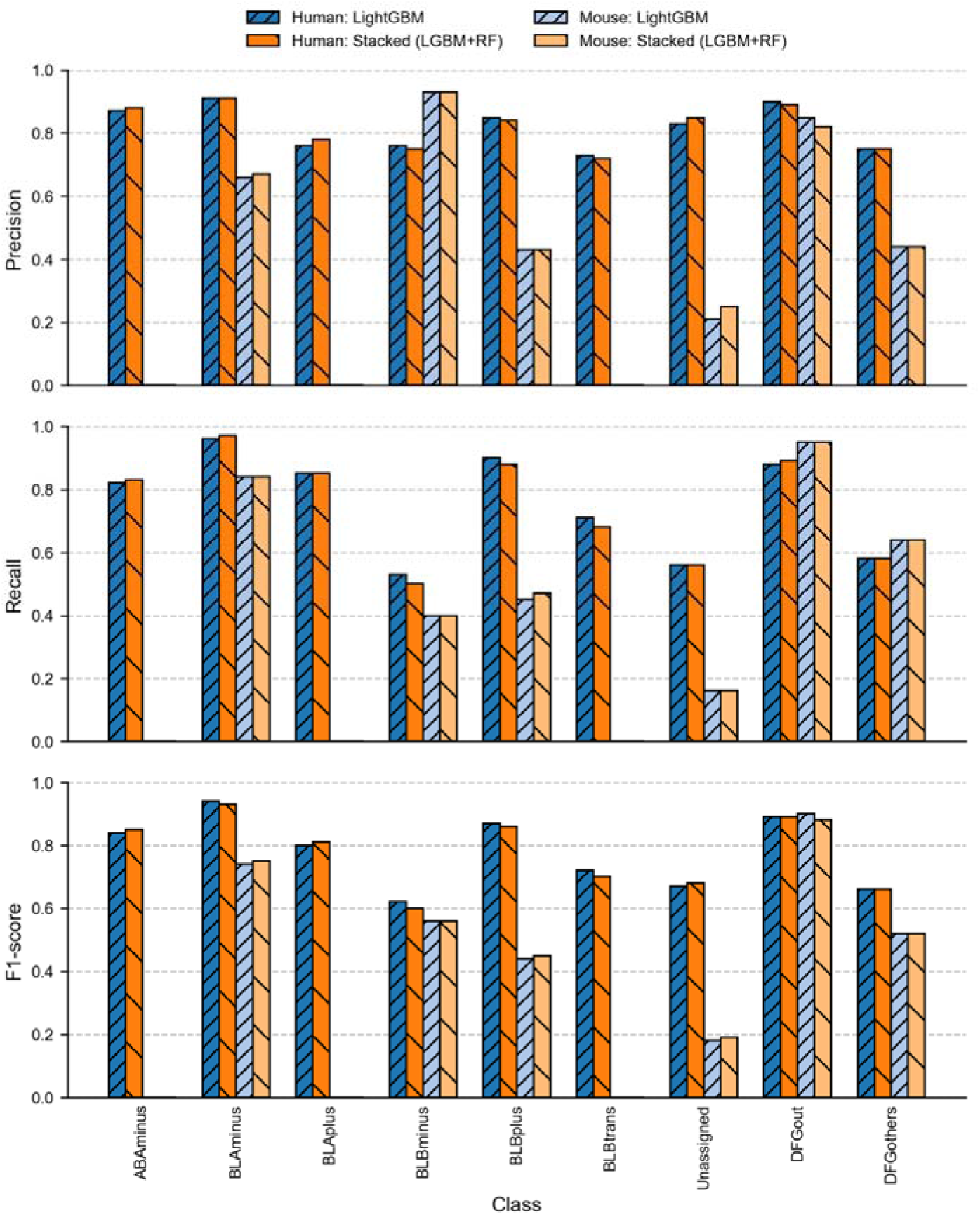
Comparison of class-wise prediction performance between LightGBM and a stacking model of LightGBM and RF. Performance is reported in terms of precision, recall, and F1-score for each class.

### 3.2 Benchmarking KASSPer Prediction for Cognate Docking

In order to examine the potential of KASSPer prediction to inform the optimal structure for SBVS, a cognate docking experiment was conducted on 25 DUD-E kinases. The pairs of kinase sequence and the SMILES string of the reference structures in the DUD-E set were utilized as inputs for KASSPer. The ligand was docked to the KASSPer predicted state. To provide a basis for comparison, ensemble docking was performed using MSM models. The compound was docked to all predicted MSM models, and the conformation with the lowest predicted binding affinity was selected for analysis. The individual result is given in Supplementary data Table S4 and summarized in Table 2.

**Table 2.** Cognate docking results of 25 kinases using KASSPer prediction and ensemble docking. For ensemble docking, the kinase conformation with the lowest binding affinity was selected for evaluation. The docking success is defined as ligand RMSD less than or equal to 2 Å.

| Method | Conformation Prediction Success | Average RMSD | Docking Success |
| --- | --- | --- | --- |
| KASSPer | 92% | 3.518 Å | 56% |
| Ensemble Docking | 44% | 4.417 Å | 32% |

KASSPer demonstrated a high degree of accuracy in predicting the conformational state of the reference complex structure (92%), while the docking score proved ineffective in guiding the correct kinase conformational state, with a success rate of 44%. In a similar vein, the RMSD and the success rate of the KASSPer prediction are both better than those of the ensemble docking method. Accurate prediction of the receptor conformational state had a significant impact on docking performance. In particular, it improved the accuracy of predicted binding poses. The two KASSPer cases that did not achieve state prediction success were EGFR and IGF1R, with RMSDs of 8.433 Å and 7.020 Å, respectively. In the context of ensemble docking, the 11 cases that exhibited the correct conformational state demonstrated an average RMSD of 2.714 Å, while the cases that did not meet this standard exhibited an average RMSD of 5.756 Å, implying that finding a proper protein conformation is important for predicting docking pose precisely.

BRAF serves as an example, illustrating the success of KASSPer prediction to obtain the conformational state and docking pose. In contrast, ensemble docking demonstrated a failure to predict both. The reference complex structure has been classified as DFGin_BLAminus state by KinCoRe. KASSPer’s prediction of the conformational state was accurate, and the docking pose exhibited an RMSD of 1.384 Å (Figure 2A). However, the lowest AutoDock score was obtained for DFGout conformation, yielding an RMSD of 7.260 Å (Figure 2B).

**Figure 2.**
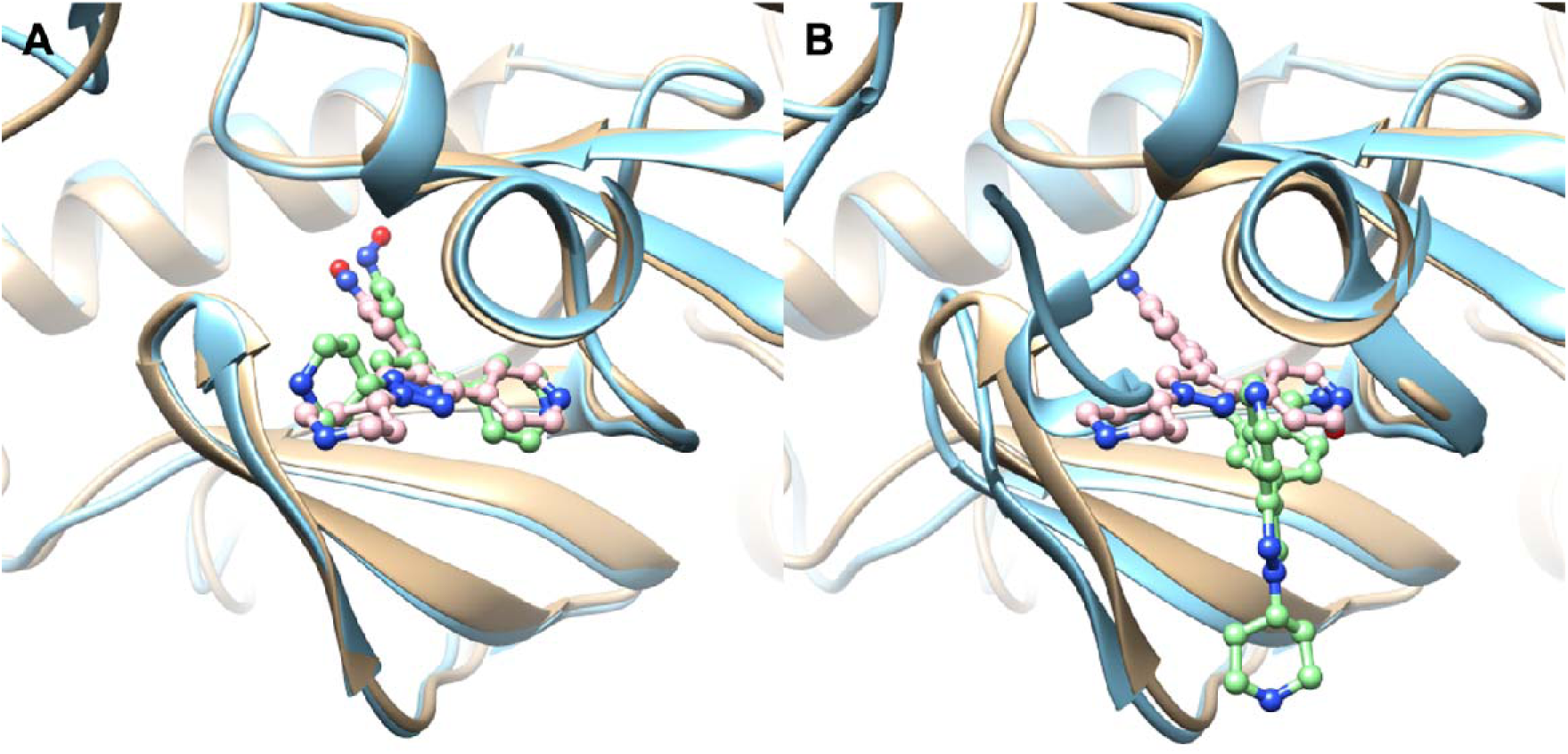
Predicted poses of BRAF cognate docking experiment. Reference structure (PDB ID: 3D4Q) is superimposed on the predicted structures. The cognate ligand docked to (**A**) KASSPer predicted structure, DFGin_BLAminus, and (**B**) the lowest AutoDock score, DFGout_BBAminus. The crystal and predicted protein structures are colored gold and sky blue, respectively, while the ligands have pink and green colors.

Overall, these results indicate that accurate prediction of kinase conformational state is associated with improved docking pose prediction. By providing a reasonable estimate of the ligand-competent receptor conformation, KASSPer may help reduce uncertainty in docking calculations. This observation indicates that KASSPer predictions could be useful as an initial receptor model in SBVS of kinases with substantial conformational variability.

### 3.3 Applying KASSPer to SBVS

As the next step, we utilized KASSPer prediction for kinase SBVS. For KASSPer screening, the compounds were docked to the KASSPer-predicted conformation and ranked by AutoDock scores. On the other hand, ensemble screening was performed by docking the molecules to all kinase conformations generated by MSM. The lowest predicted binding affinity was selected as a representative score for ranking the compound. A comparison of the two methods is presented in Table 3, and detailed results for individual targets are also provided in Supplementary data S5.

**Table 3.**
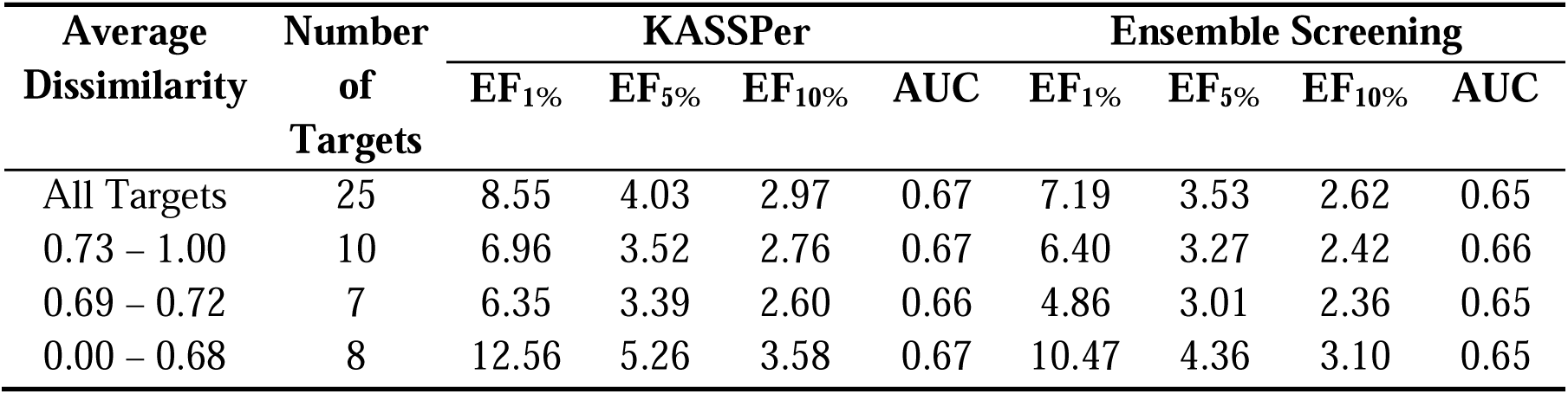
Virtual screening benchmarks of kinases using KASSPer prediction and ensemble screening. The targets are classified by the average dissimilarity of active compounds.

| Average<br>Dissimilarity | Number<br>of<br>Targets | KASSPer |  |  |  | Ensemble Screening |  |  |  |
| --- | --- | --- | --- | --- | --- | --- | --- | --- | --- |
|  |  | EF<sub>1%</sub> | EF<sub>5%</sub> | EF<sub>10%</sub> | AUC | EF<sub>1%</sub> | EF<sub>5%</sub> | EF<sub>10%</sub> | AUC |
| All Targets | 25 | 8.55 | 4.03 | 2.97 | 0.67 | 7.19 | 3.53 | 2.62 | 0.65 |
| 0.73 – 1.00 | 10 | 6.96 | 3.52 | 2.76 | 0.67 | 6.40 | 3.27 | 2.42 | 0.66 |
| 0.69 – 0.72 | 7 | 6.35 | 3.39 | 2.60 | 0.66 | 4.86 | 3.01 | 2.36 | 0.65 |
| 0.00 – 0.68 | 8 | 12.56 | 5.26 | 3.58 | 0.67 | 10.47 | 4.36 | 3.10 | 0.65 |

On average, KASSPer screening exhibited superior performance in all metrics when compared with ensemble screening. For EF_1%_, KASSPer (8.55) exhibited an 18.9% improvement over the ensemble method (7.19). The statistical analysis was conducted using the paired t-test, yielding p-values less than 0.05 (0.0225). Similarly, KASSPer also improved in both EF_5%_ (14.2%) and EF_10%_ (13.4%), and showed statistical significance with p-values less than 0.05, 0.0211 and 0.0152 for EF5% and EF10%, respectively.

The utilization of multiple conformations of the target protein in the ensemble screening technique is advantageous for the identification of diverse hit molecules. Song et al. (Song *et al*. 2024) calculated the dissimilarity of active compounds of the DUD-E kinase subset using RDKit (Landrum *et al*. 2025) and found that the ensemble screening using MSM models outperformed the single conformation screening using AF2 and AF3 structures, especially for the targets with high dissimilarity. To investigate whether a screening against the conformer with KASSPer prediction still works for diverse active compounds, we classified the targets into three groups by the average dissimilarity of active compounds (Table 3). Across all target classes and metrics, KASSPer demonstrated superior performance in comparison to the ensemble screening.

One of the examples that KASSPer-based SBVS outperformed ensemble screening is PRKCB. The EF_1%_ of KASSPer screening and ensemble screening are 21.55 and 12.63, respectively. Figure 3 shows the docking poses of one of the active compounds, CHEMBL6291. KASSPer predicted the compound binds to DFGin_BLAminus, resulting a docking score as -11.31 kcal/mol (Figure 3A). The KASSPer-based screening ranked the compound within the top 1% (38^th^) out of 8827 compounds. On the other hand, by the ensemble screening, the lowest binding energy was obtained from the Unassigned_Unassigned conformation (-11.45 kcal/mol, Figure 3B). Although the ensemble screening found the lower energy conformation, it did not rank the molecule within the top 1% (111^th^)

**Figure 3.**
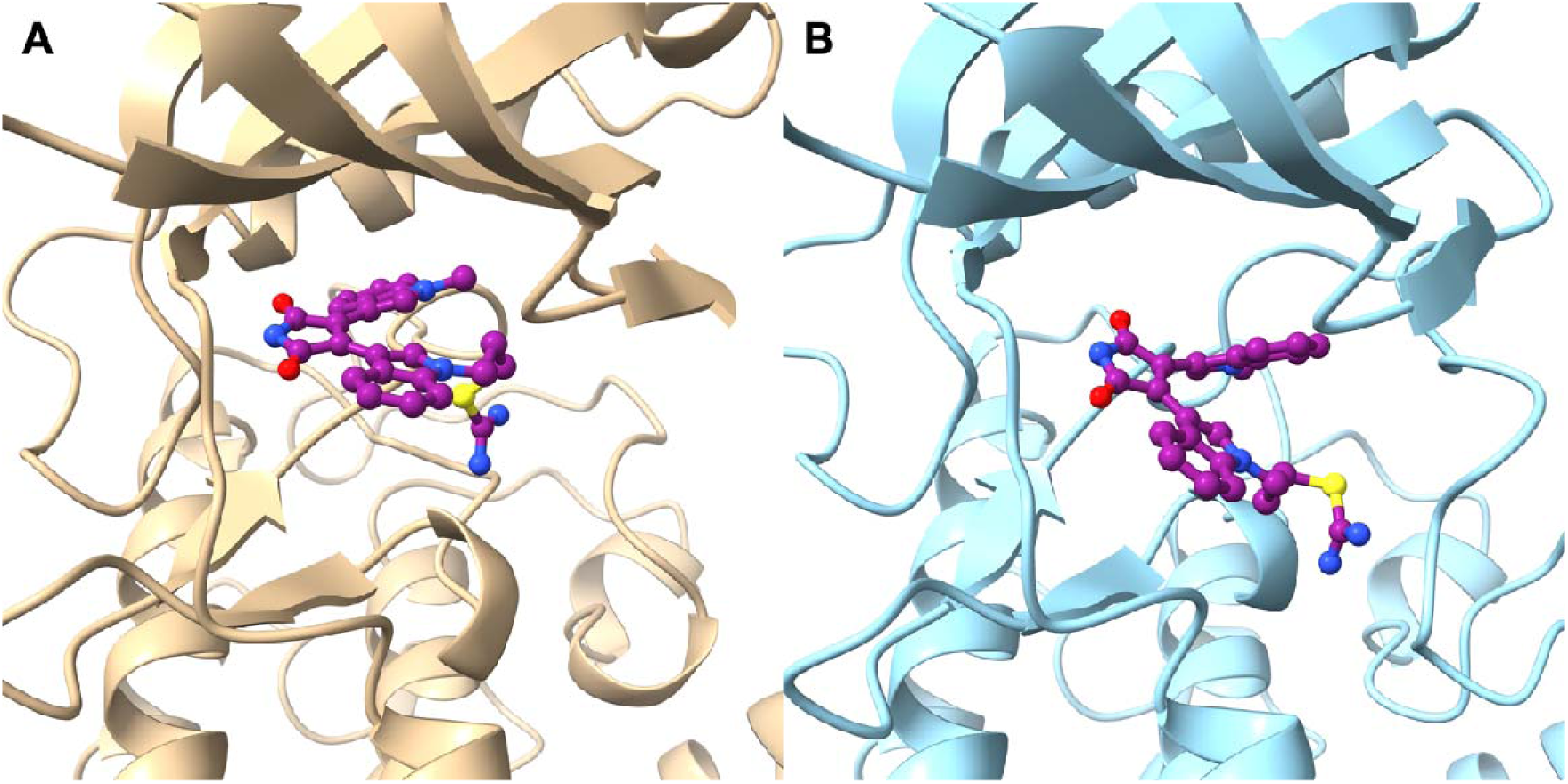
Predicted binding poses of CHEMBL6291, an active compound of PRKCB, to (**A**) KASSPer predicted structure, DFGin_BLAminus (gold), and (**B**) the Unassigned (sky blue) with the lowest AutoDock score. The docked ligand is colored purple in both panels.

In summary, these results suggest that KASSPer-based virtual screening can achieve better performance than ensemble-based screening for kinases. Across the DUD-E kinase subset, screening against a single KASSPer-predicted conformation yielded consistently higher enrichment metrics than ensemble screening, including for targets with high ligand dissimilarity. These findings support the use of KASSPer for kinase SBVS. The method reduces the number of docking calculations required for ensemble screening while maintaining accuracy.

## 4. Conclusion

A significant challenge in the field of SBVS is the flexibility of receptors, with kinases serving as a typical example. These enzymes exhibit a high degree of conformational flexibility at their active site, depending on the bound ligand. While the ensemble screening may offer a solution to this issue, it should be noted that the number of computations increases in proportion to the number of conformations. To address this issue, we developed KASSPer, a language model-based framework to predict kinase conformation when a compound binds. A stacking model of LightGBM and RF was selected based on the validation result. The program demonstrated a high degree of accuracy in predicting the kinase conformation for the cognate docking benchmark, thereby facilitating the precise prediction of docking poses. In the context of the SBVS benchmark, KASSPer demonstrated superior performance in comparison to ensemble screening, as indicated by EF_1%_, EF_5%_, EF_10%_, and AUC. The findings indicate that KASSPer exhibits superiority in capturing ligand-induced kinase state variability in comparison to conventional ensemble methods.

Despite strong screening performance, the embeddings used by KASSPER are not directly interpretable, which hinders biological insight into sequence contributions to DFG state selection. Future work will focus on designing embedding strategies with built-in feature attribution to reveal which protein residues and ligand substructures most strongly influence state predictions. Such insights will enhance model interpretability and guide rational design of kinase modulators. Overall, KASSPer offers a new paradigm for efficient and interpretable virtual screening of kinase targets.

## Author Contributions

Wonkyeong Jang (Formal analysis [lead], Data Curation [lead], Software [lead], Writing – original draft [lead]), Woong-Hee Shin (Conceptualization [lead], Project administration [lead], Supervision [lead], Writing – original draft [supporting], Writing – review & editing [lead])

## Supplementary Data

Supplementary data is available at Bioinformatics online. Conflict of interest: None declared.

## Data Availability

The receptor structures predicted by MSM protocol are available at https://zenodo.org/records/8272608. DUD-E benchmark compound library is available at dude.docking.org.

## Code Availability

The source code for KASSPer is available at https://github.com/kucm-lsbi/KASSPer.

