## Supplementary data for "KASSPer: Kinase Active Site Structure Prediction using Protein and Ligand Language Models and Its Application to Virtual Screening"

**Table S1**. Hyperparameters for the machine learning models benchmarked.

| Models | Name | Parameters |
|---|---|---|
| XGBoost | n_estimators | 50, 100, 200 |
| | max_depth | 3, 5, 7 |
| | learning_rate | 0.01, 0.1, 0.2 |
| | colsample_bytree | 0.7, 1.0 |
| LightGBM | n_estimators | 50, 100, 200 |
| | max_depth | 3, 5, 7 |
| | learning_rate | 0.01, 0.1, 0.2 |
| | bagging_fraction | 0.7, 1.0 |
| | feature_fraction | 0.7, 1.0 |
| CatBoost | tuned iterations | 50, 100, 200 |
| | max_depth | 3, 5, 7 |
| | learning_rate | 0.01, 0.1, 0.2 |
| | subsample | 0.7, 1.0 |
| | rsm | 0.7, 1.0 |
| Random Forest | n_estimators | 50, 100, 200 |
| | max_depth | None, 10, 20 |
| | max_features | sqrt, log2 |
| | min_samples_split | 2, 5 |
| Logistic Regression | hyperparameter C | 0.01, 0.1, 1, 10 |
| | max_iter | 100, 200 |
| | multi_class | ovr, multinomial |
| | class_weight | None, balanced |
| SVM | hyperparameter C | 0.01, 0.1, 1, 10 |
| | kernel function | linear, rbf |
| | gamma | scale, auto |
| | max_iter | 100, 200 |

**Table S2.** Performance of individual models.

| Sets | Single Model | | | | | | | | Stacking Ensemble | | |
|---|---|---|---|---|---|---|---|---|---|---|---|
| | **XG Boost** | **Light GBM** | **Cat Boost** | **Random Forest** | **Logistic Regression** | **SVM** | **Cross Attention** | **Concatenate** | **XGBoost + LightGBM** | **Light GBM + Random Forest** | **XGBoost + Random Forest** |
| Human | | | | | | | | | | | |
| MCC | 0.80 | 0.82 | 0.77 | 0.80 | 0.73 | 0.51 | 0.77 | 0.74 | 0.81 | 0.82 | 0.81 |
| Precision | 0.79 | 0.82 | 0.76 | 0.79 | 0.72 | 0.59 | 0.72 | 0.72 | 0.82 | 0.82 | 0.80 |
| Recall | 0.73 | 0.75 | 0.69 | 0.74 | 0.65 | 0.56 | 0.68 | 0.67 | 0.74 | 0.75 | 0.74 |
| F1-Score | 0.75 | 0.78 | 0.71 | 0.75 | 0.67 | 0.56 | 0.70 | 0.69 | 0.77 | 0.78 | 0.77 |
| Mouse | | | | | | | | | | | |
| MCC | 0.60 | 0.55 | 0.53 | 0.58 | 0.41 | 0.29 | 0.51 | 0.45 | 0.56 | 0.56 | 0.59 |
| Precision | 0.45 | 0.44 | 0.43 | 0.46 | 0.31 | 0.27 | 0.40 | 0.37 | 0.45 | 0.44 | 0.44 |
| Recall | 0.46 | 0.43 | 0.38 | 0.41 | 0.31 | 0.27 | 0.41 | 0.51 | 0.43 | 0.43 | 0.45 |
| F1-Score | 0.75 | 0.78 | 0.71 | 0.75 | 0.67 | 0.56 | 0.70 | 0.69 | 0.77 | 0.78 | 0.77 |

**Table S3.** Detailed results of LightGBM and a stacking model of LightGBM with Random Forest for the individual class. For the mouse set, no crystal structures was assigned as BLBtrans.

| Conformational State | Human | | Mouse | |
|---|---|---|---|---|
| | **LightGBM** | **Stacking** | **LightGBM** | **Stacking** |
| | | **Precision** | | |
| ABAminus | 0.87 | 0.88 | 0.00 | 0.00 |
| BLAminus | 0.91 | 0.91 | 0.66 | 0.75 |
| BLAplus | 0.76 | 0.78 | 0.00 | 0.00 |
| BLBminus | 0.76 | 0.75 | 0.93 | 0.92 |
| BLBplus | 0.85 | 0.84 | 0.43 | 0.42 |
| BLBtrans | 0.73 | 0.72 | 0.00 | 0.00 |
| Unassigned | 0.83 | 0.85 | 0.21 | 0.21 |
| DFGout | 0.90 | 0.89 | 0.85 | 0.80 |
| DFGothers | 0.75 | 0.75 | 0.44 | 0.44 |
| | | **Recall** | | |
| ABAminus | 0.82 | 0.82 | 0.00 | 0.00 |
| BLAminus | 0.96 | 0.96 | 0.84 | 0.86 |
| BLAplus | 0.85 | 0.85 | 0.00 | 0.00 |
| BLBminus | 0.53 | 0.50 | 0.40 | 0.56 |
| BLBplus | 0.9 | 0.85 | 0.45 | 0.47 |
| BLBtrans | 0.71 | 0.68 | 0.00 | 0.00 |
| Unassigned | 0.56 | 0.58 | 0.16 | 0.16 |
| DFGout | 0.88 | 0.86 | 0.95 | 0.87 |
| DFGothers | 0.58 | 0.56 | 0.64 | 0.64 |
| | | **F1-Score** | | |
| ABAminus | 0.84 | 0.86 | 0.00 | 0.00 |
| BLAminus | 0.94 | 0.93 | 0.74 | 0.80 |
| BLAplus | 0.8 | 0.82 | 0.00 | 0.00 |
| BLBminus | 0.62 | 0.61 | 0.56 | 0.69 |
| BLBplus | 0.87 | 0.84 | 0.44 | 0.44 |
| BLBtrans | 0.72 | 0.70 | 0.00 | 0.00 |
| Unassigned | 0.67 | 0.66 | 0.18 | 0.18 |
| DFGout | 0.89 | 0.86 | 0.90 | 0.83 |
| DFGothers | 0.66 | 0.61 | 0.52 | 0.52 |

**Table S4.** Individual result of the cognate docking experiment. The structural states of the ensemble docking were selected based on the lowest AutoDock-GPU score.

| Protein | Conformational State | KASSPer | | Ensemble Docking | |
|---|---|---|---|---|---|
| | | Predicted State | RMSD (Å) | Lowest Scoring State | RMSD (Å) |
| ABL1 | DFGout | DFGout | 0.293 | DFGout | 0.293 |
| AKT1 | BLAminus | BLAminus | 0.574 | BLAminus | 0.574 |
| AKT2 | BLAminus | BLAminus | 0.719 | BLAminus | 0.719 |
| BRAF | BLAminus | BLAminus | 1.384 | DFGout | 7.260 |
| CDK2 | BLBtrans | BLBtrans | 9.652 | DFGout | 11.369 |
| CSF1R | DFGout | DFGout | 8.210 | BLBtrans | 7.819 |
| EGFR | BLBplus | BLAminus | 8.433 | BLBtrans | 10.028 |
| FGFR1 | BLAplus | BLAplus | 0.347 | DFGout | 0.890 |
| IGF1R | DFGothers | Unassigned | 7.020 | DFGout | 6.904 |
| JAK2 | ABAminus | ABAminus | 1.317 | DFGout | 6.925 |
| KDR | DFGout | DFGout | 1.234 | DFGout | 1.234 |
| KIT | DFGout | DFGout | 4.902 | DFGout | 4.902 |
| LCK | BLAminus | BLAminus | 0.734 | BLBplus | 6.658 |
| MAP2K1 | BLBplus | BLBplus | 2.589 | BLBplus | 2.589 |
| MAPK1 | BLAminus | BLAminus | 1.205 | DFGothers | 3.456 |
| MAPK10 | ABAminus | ABAminus | 9.173 | BLAplus | 8.842 |
| MAPK14 | BLBplus | BLBplus | 6.711 | BLAminus | 2.587 |
| MAPKAPK2 | BLAminus | BLAminus | 0.819 | BLBplus | 1.538 |
| MET | DFGout | DFGout | 11.334 | DFGout | 11.334 |
| PLK1 | BLAminus | BLAminus | 1.856 | Unassigned | 3.280 |
| PRKCB | BLAminus | BLAminus | 4.349 | BLAminus | 4.349 |
| PTK2 | Unassigned | Unassigned | 0.881 | Unassigned | 0.881 |
| ROCK1 | BLAminus | BLAminus | 1.223 | DFGout | 3.021 |
| TGFBR1 | BLAminus | BLAminus | 0.881 | BLAminus | 0.881 |
| WEE | BLAminus | BLAminus | 2.100 | BLAminus | 2.100 |

**Table S5.** Virtual screening benchmark result of KASSPer and ensemble docking for DUD-E kinase subset.

| **Kinase** | **Average Dissimilarity** | **KASSPer** | | | | **Ensemble Screening** | | | |
|---|---|---|---|---|---|---|---|---|---|
| | | EF1% | EF5% | EF10% | AUC | EF1% | EF5% | EF10% | AUC |
| ABL1 | 0.73 | 10.47 | 4.62 | 3.30 | 0.71 | 9.36 | 3.74 | 2.47 | 0.64 |
| AKT1 | 0.68 | 9.57 | 5.19 | 3.92 | 0.74 | 12.64 | 5.46 | 3.79 | 0.74 |
| AKT2 | 0.69 | 16.26 | 5.82 | 3.76 | 0.77 | 12.84 | 5.65 | 3.93 | 0.78 |
| BRAF | 0.70 | 9.86 | 4.08 | 3.02 | 0.72 | 7.23 | 5.26 | 3.22 | 0.71 |
| CDK2 | 0.78 | 3.16 | 1.73 | 1.53 | 0.61 | 5.70 | 2.57 | 2.19 | 0.66 |
| CSF1R | 0.73 | 6.03 | 3.13 | 2.71 | 0.63 | 5.43 | 3.50 | 2.17 | 0.64 |
| EGFR | 0.70 | 6.45 | 3.84 | 3.01 | 0.65 | 4.42 | 3.06 | 2.21 | 0.60 |
| FGFR1 | 0.69 | 6.50 | 5.03 | 3.81 | 0.70 | 2.16 | 2.45 | 2.16 | 0.66 |
| IGF1R | 0.69 | 4.75 | 3.38 | 2.84 | 0.72 | 6.78 | 3.24 | 2.97 | 0.74 |
| JAK2 | 0.73 | 5.61 | 2.43 | 1.68 | 0.57 | 5.61 | 2.43 | 1.87 | 0.58 |
| KDR | 0.75 | 9.30 | 4.60 | 3.35 | 0.70 | 9.06 | 4.60 | 3.40 | 0.70 |
| KIT | 0.73 | 12.67 | 5.06 | 3.92 | 0.72 | 12.67 | 4.09 | 3.01 | 0.70 |
| LCK | 0.73 | 6.19 | 3.19 | 2.52 | 0.66 | 4.28 | 2.90 | 2.33 | 0.62 |
| MAP2K1 | 0.71 | 0.00 | 0.99 | 0.91 | 0.58 | 0.00 | 0.83 | 0.83 | 0.56 |
| MAPK1 | 0.75 | 11.46 | 5.58 | 3.80 | 0.74 | 6.37 | 4.82 | 3.67 | 0.72 |
| MAPK10 | 0.58 | 2.89 | 2.31 | 2.50 | 0.66 | 3.85 | 2.89 | 2.31 | 0.65 |
| MAPK14 | 0.75 | 1.90 | 1.94 | 1.73 | 0.64 | 3.46 | 2.49 | 1.90 | 0.64 |
| MAPKAPK2 | 0.68 | 5.99 | 3.77 | 3.17 | 0.68 | 3.99 | 2.78 | 2.67 | 0.67 |
| MET | 0.70 | 0.60 | 0.60 | 0.84 | 0.48 | 0.60 | 0.60 | 1.20 | 0.52 |
| PLK1 | 0.65 | 6.55 | 4.30 | 3.37 | 0.62 | 1.87 | 1.50 | 1.68 | 0.54 |
| PRKCB | 0.56 | 21.55 | 7.86 | 5.04 | 0.64 | 12.63 | 5.49 | 3.78 | 0.67 |
| PTK2 | 0.68 | 0.00 | 0.60 | 0.60 | 0.46 | 0.00 | 0.00 | 0.40 | 0.37 |
| ROCK1 | 0.74 | 2.00 | 2.60 | 2.80 | 0.72 | 0.00 | 1.20 | 1.10 | 0.68 |
| TGBFR1 | 0.64 | 5.28 | 4.36 | 3.01 | 0.69 | 5.28 | 3.60 | 3.31 | 0.69 |
| WEE1 | 0.39 | 48.64 | 13.71 | 7.06 | 0.89 | 43.48 | 13.16 | 6.86 | 0.88 |
| Average | | 8.55 | 4.03 | 2.97 | 0.67 | 7.19 | 3.53 | 2.62 | 0.65 |

**Figure S1.** The class distribution of kinase crystal structures. The y-axis is the percentage of the class.

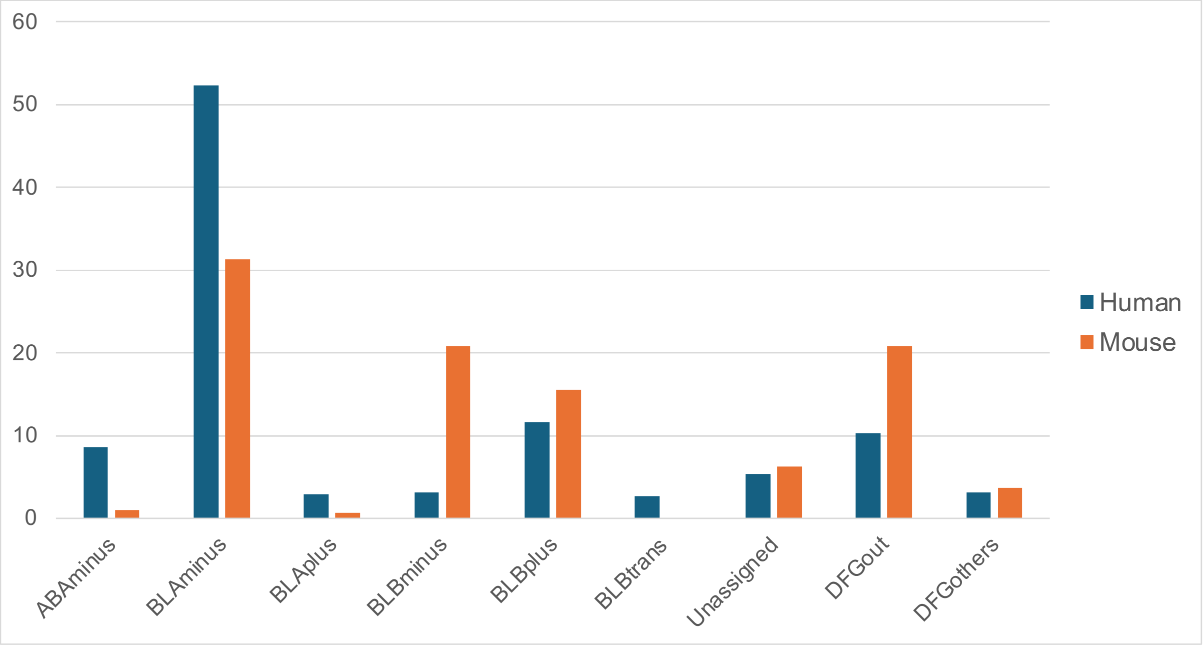


**Figure S2.** Analysis between the human test set and the mouse set. **A**. Sequence identity distribution. **B.** Tanimoto similarity distribution.

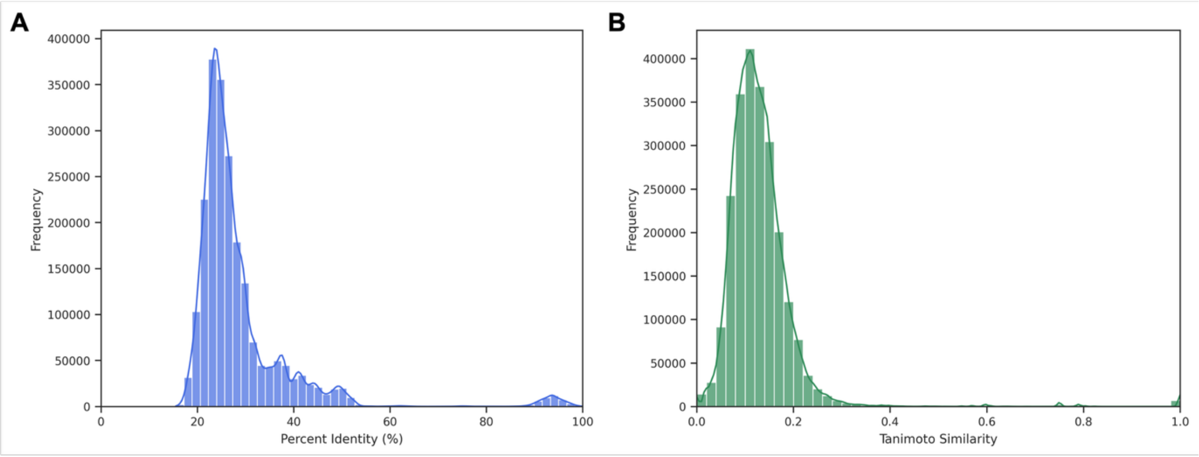
